# A fish rots from the head down: how to use the leading digits of ecological data to detect their falsification

**DOI:** 10.1101/368951

**Authors:** J. Cerri

**Affiliations:** Institute of Management, Scuola Superiore Sant’Anna, Piazza Martiri della Libertà 33, 56127, Pisa, Italy; Istituto Superiore per la Protezione e la Ricerca Ambientale, ISPRA

## Abstract

Managing wildlife populations requires good data. Researchers and policy makers need reliable population estimates and, in case of commercial or recreational harvesting, also trustworthy information about the number of removed individuals. However, auditing schemes are often weak and political or economic pressure could lead to data fabrication or falsification. Time-series data and population models are crucial to detect anomalies, but they are not always available nor feasible. Therefore, researchers need other tools to identify suspicious patterns in ecological and environmental data, to prioritize their controls. We showed how the Benford’s law might be used to identify anomalies and potential manipulation in ecological data, by testing for the goodness-of-fit of the leading digits with the Benford’s distribution. For this task, we inspected two datasets that were found to be falsified, containing data about estimated large carnivore populations in Romania and Soviet commercial whale catches in the Pacific Ocean. In both the two datasets, the first and second digits numerical series deviated from the expected Benford’s distribution. In data about large carnivores, the first too digits, taken together, also deviated from the expected Benford’s distribution and were characterized by a high Mean Absolute Deviation. In Soviet whale catches, while the single digits deviated from the Benford’s distribution and the Mean Absolute Deviation was high, the first two digits were not anomalous. This controversy invites researchers to combine multiple measures of nonconformity and to be cautious in analyzing mixtures of data. Testing the distribution of the leading digits might be a very useful tool to inspect ecological datasets and to detect potential falsifications, with great implications for policymakers and researchers as well. For example, if policymakers revealed anomalies in harvesting data or population estimates, commercial or recreational harvesting could be suspended and controls strengthened. On the other hand, revealing falsification in ecological research would be crucial for evidence-based conservation, as well as for research evaluation.

## Introduction

Successful management of animal and plant populations requires informed decision-making. Information about populations and their geographical distribution is crucial for designing effective networks of protected areas, identifying threats and integrating conservation in policy making. Furthermore, as many animals and plants are traded, environmental managers also need trustworthy information about the number and qualities of these individuals which are removed from nature. During the last 20 years, conservation biology was flooded with information. Digitalization enabled conservationists and agencies to store and share their data (Hampton et al., 2013; Page et al., 2015). Advances in informatics and the computational power of computers, allowed for an unprecedented large-scale adoption of statistics in environmental management and nowadays data analysis and scientific evidence are the prerequisite for many conservation policies worldwide (Dubois et al., 2017). However, the debate about data quality was partial somehow. Unreliable ecological information was believed to stem from inadequate monitoring or superficial statistical inference and modeling (Legg and Nagy, 2006; Sutherland, 2006), while other elephants in the room, like data manipulation or fabrication, went relatively unnoticed. While this topic certainly makes most scientists and practitioners uncomfortable, there are some good reasons to believe that some ecological and environmental data get sanitized, manipulated or deliberately fabricated. The first reason is the unprecedented commercial pressure affecting many animal and plant species. Wildlife commerce is one of the largest worldwide (Symes et al., 2018), and the demand for specific animal or plant based products changes relentlessly due to fads (https://www.theguardian.com/environment/2018/apr/27/stolen-succulents-california-hipster-plants-at-center-of-smuggling-crisis) or complex socio-economic dynamics (Duffy and St. John, 2013; Duffy et al., 2016). This, in turns, can generate considerable political pressures over those researchers who are responsible for ecological census or harvesting quotas (Darimont et al., 2018). In the absence of effective control schemes and stewardship norms, the consequences of these pressures can be disastrous. In 2017 the Romanian government halted the recreational hunting of large carnivores, after that growth rates of the bear population, a valuable game, were found out to be biologically unrealistic and prone to falsification (Popescu et al., 2016). Again, retrospective analysis demonstrated that in the Soviet Union whaling data were misreported for decades, due to the perverse economic incentives introduced by unrealistic economic targets (Clapham and Ivashchenko, 2009; Ivashchenko et al., 2011, 2013). The second reason lies in the fact that some researchers manipulate or falsify their data to obtain the desired outcomes (Fanelli, 2009). Scientific misconduct is a plague in many disciplines adopting easily falsifiable data collection modes, such as questionnaires or laboratory experiments. Although ecology and environmental sciences are characterized by more time-consuming and collective methods of research, which are likely to discourage lone wolves and to promote whistleblowing (Barlow et al., 2018), ecology experienced the same changes in funding and tenuring that characterized other disciplines: an overall reduction of resources coupled with an extinction of long-term funding (Bakker et al., 2010; Kuebbing et al., 2018) and the imposition of ‘publish-or-perish’ policies. These changes inevitably lead to scientific misconduct (Grimes et al., 2018). Finally, the reduction of financial resources for research in conservation, coupled with the ongoing economic crisis, might also encourage the large-scale replacement of professionals with volunteers (Lewandowski and Specht, 2015), which sometimes have serious conflicts of interest making them prone to sanitize their data. This mix of economic pressures, shortsighted research funding and voluntary engagement is too dangerous to be ignored. While scientific misconduct can be reduced through long-term changes, like the enforcement of control mechanisms, or the promotion of research integrity, we believe that short-term responses are needed too. It is time for ecologists and conservationists to start scrutinizing the quality of available data, and to prioritize the inspection of those that look suspicious. The detection of manipulated or fabricated data has received considerable attention in the last few years, across many different sectors (e.g. finance,Michalski and Stolz, 2013; Rauch, Göttsche and Langenegger, 2014; e.g. political sciences, Beber and Scacco, 2012; Mebane, 2008; e.g. physics Brumfiel, 2002) and various approaches are now available. This research wants to encourage their use in ecology and conservation, by showing how relatively simple statistical tests for numerical digits might indicate anomalies in ecological datasets. We will use two datasets which were found to be manipulated, as a validated case study.

## Materials and methods

### Statistical detection of manipulated data

The statistical detection of falsified data includes both supervised and unsupervised approaches (Bolton and Hand, 2002). Supervised techniques require some prior knowledge to classify observations as true or frauds (e.g neural networks,Hodge and Austin, 2004), or to develop a theoretical model generating those data that are expected to occur, absent fraud, to compare them with real ones (Popescu et al., 2016). Unsupervised techniques do not require any particular prior knowledge and test if observed data significantly depart from some sort of expected values. Some unsupervised approaches for example exploit cognitive bias affecting number generation by humans (Beber and Scacco, 2012; Klimek et al., 2012; Kobak, Shpilkin and Pshenichnikov, 2016; Nigrini, 2012; Pitt and Hill, 2013), or test whether reported statistics are compatible with the granularity of the data (Anaya, 2016). Digit tests based on the Benford’s law are the most common unsupervised approach. In 1938, Frank Benford (Benford, 1938)observed that first and second digits in numerical series follow a particular logarithmic distribution (Eq. 1), as it had been previously suggested byNewcomb (1881). Since then, various natural phenomena have been found to follow this distribution (Campos, Salvo and Flores-Moya, 2016; Sambridge, Tkalčić and Jackson, 2010). In statistical fraud detection, empirical first and second digits of inspected data are compared with the Benford’s distribution and if they show significant departures, data are generally deemed to require further investigations (Durtschi, Hillison and Pacini, 2004; Nigrini, 1996, 2012). Absent fabrication, the first and second digits of any numerical series follow the Benford’s distribution, provided that: sample size is greater than 100, the data measure the same concept, the data are not numbers that have been allocated a-priori (e.g. identification numbers), data distribution is skewed to the left, with the mean greater than the median and data are not too clustered around the mean (Durtschi et al. 2004; Fewster, 2009; Hill, 1995 a,b,c;Leemis, Schmeiser and Evans, 2000). The Benford’s law is scale-invariant and base-invariant, so even transforming the data, for example by shifting from observed animals to densities, does not mask their departure from the Benford’s distribution (Hill, 1995 b,c). Typically, data conformity with the Benford’s distribution is tested with simple statistical goodness-of-fit tests, like the chi-square test (Nigrini, 2012). To date, digits conformance with the Benford’s distribution was tested to audit data in financial accountability (Nigrini, 1996, 2012), environmental chemistry (De Marchi and Hamilton, 2006), political elections (Mebane, 2011), surveys (Judge and Schechter, 2009) and statistics (Diekmann, 2007). To the best of our knowledge, the only conservation studies adopting these methods were about fisheries (Graham, Hasseldine and Paton, 2009; Tsagbey, De Carvalho and Page, 2017). A complete website containing information about the Benford’s law, altogether with some examples from the real world and a list of scientific publications, is available at http://www.benfordonline.net

### Case studies and statistical analysis

To demonstrate the potential of the Benford’s law for detecting anomalies in ecological data, we considered two datasets which were found to be manipulated. The first one was published in Popescu et al. (2016). It contained all the regional population estimates, developed by the Romanian government between 2005 and 2012, about three species of large carnivores: the brown bear (Ursus arctos), the Eurasian lynx (Lynx lynx) and the gray wolf (Canis lupus). Population growth rates of the brown bear were found to be over-optimistic, being much higher than reported growth rates from existing literature. Moreover, the difference between reported and plausible estimates showed a positive correlation with hunting pressure. On the other hand, population growth rates of the Eurasian lynx were almost entirely below the range of potential values obtained from previous studies. This might indicate that existing snow-tracking schemes for monitoring lynx might be inadequate to obtain reliable population estimates. Finally, despite being generally in line with theoretical expectations, in a few counties population growth rates of the gray wolf were above their expected values. Again, this might indicate the existence of data manipulation at the local level, although as not as widespread as for the brown bear. In this case, we tested whether the first and second digits of regional population estimates of each species, followed the Benford’s distribution or not. As suggested by Nigrini (2012), single digits estimates were removed, and we retained values greater or equal than 10. For each species we pooled together all the data from the various years and regions, to achieve a sample size greater than 100. As a second case study, we considered reported whaling data of the former Soviet whaling fleet in the Pacific Ocean. From 1947 to 1973 the Soviet Union illegally exploited the stocks of many whale species both in the Northern and in the Southern hemisphere. This exploitation was fueled by unrealistic economic targets, coupled with strong economic bonus for whalers, that made whaling one of the most lucrative activities in the Union (Clapham and Ivashchenko, 2009; Ivashchenko et al., 2011, 2013). As a result, this whaling campaign was conducted by deliberately ignoring the quotas and regulations established by the International Whaling Committee, and it targeted animals of all ages and species. It is estimated that almost 100.000 whales, killed in the Southern hemisphere, were not reported. True catches and measures were disclosed to the IWC in the 1990s only, by some former biologists working on the vessels. In this research, we considered data of Soviet whaling fleets operating in the Pacific Ocean at that time. Our dataset was obtained by combining catches from Northern and Southern Pacific, published in Ivashchenko et al. (2013) and in Clapham et al. (2009). In this case, to achieve a suitable sample size, we pooled together the catches of five different species between 1946 and 1979: the blue whale (Balaenoptera musculus), the fin whale (Balaenoptera physalus), the humpback whale (Megaptera novaeangliae), the sei whale (Balaenoptera borealis) and the sperm whale (Physeter macrocephalus). Pooling together all the data about different species would enable to draw conclusions about the overall quality of the dataset, in this case, the former Soviet whaling system acting in the Pacific Ocean at that time. We retained catches greater, or equal, than 10. As suggested by Nigrini (2012) and Diekmann (2007), we adopted the chi-square goodness-of-fit test to check whether the first digits, the second digits, and the first couple of digits deviated from the expected Benford’s distribution. In the chi-square goodness-of-fit test, the null hypothesis states that frequencies come a Benford’s distribution: if the chi-square test was significant, we would accept the alternative hypothesis that the data do not came from this type of distribution. Therefore, a significant chi-square test would indicate some anomalous pattern in the data, that might indicate manipulation and that deserve further inspections. We also measured the Mean Absolute Deviation (MAD) of the first two digits, a robust proxy of conformity to the Benford’s distribution for two-digits series. The MAD measures the difference between absolute and expected proportions of the first couple of digits, weighted on the basis of the number of bins, equal to 90 for couples of digits. As suggested by Nigrini (2012), a value of the MAD above 0.0044 indicates non-conformity with the Benford’s distribution. The MAD index was chosen as it is relatively robust for small and large sample size. Goodness-of-fit testing and the computation of the MAD index were carried out through the statistical software “R” (RCoreTeam, 2018), with the package ‘benford.analysis’ (Cinelli, 2014).

## Results

The distributions of both large carnivore estimates and whaling data were suitable for goodness-of-fit testing: they were positively skewed, their mean was greater than the median and they had a relatively large standard deviation (Fig. 1). The distribution of the first and second digits, as well as the distribution of the first couple of digits, of large carnivore data from Popescu et al. (2016), did not conform to the Benford’s distribution. This was evident from a graphical inspection of frequency histograms, characterized by an anomalous high frequency of high digits. Moreover, for all the three species, the chi-square test indicated that neither the first digit, nor the second digit, nor the first couple of digits, conformed to a Benford’s distribution. Nonconformity of the first two digits was confirmed by the very high MAD (Fig. 2).

**Figure 1: Fig.1.**
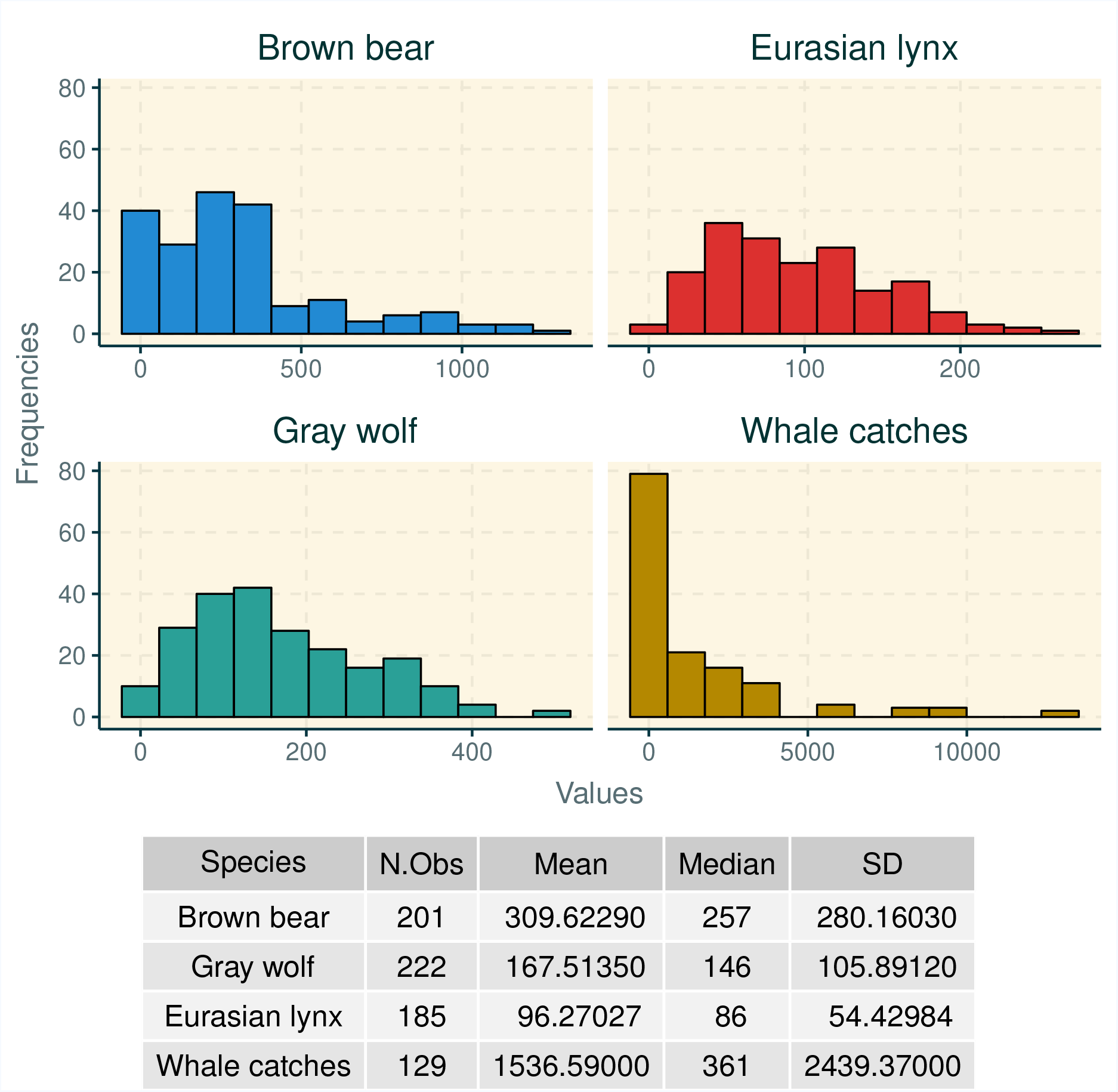
Distribution of large carnivore population data and Soviet whale catches in the Pacific Ocean.

**Figure 2: Fig.2.**
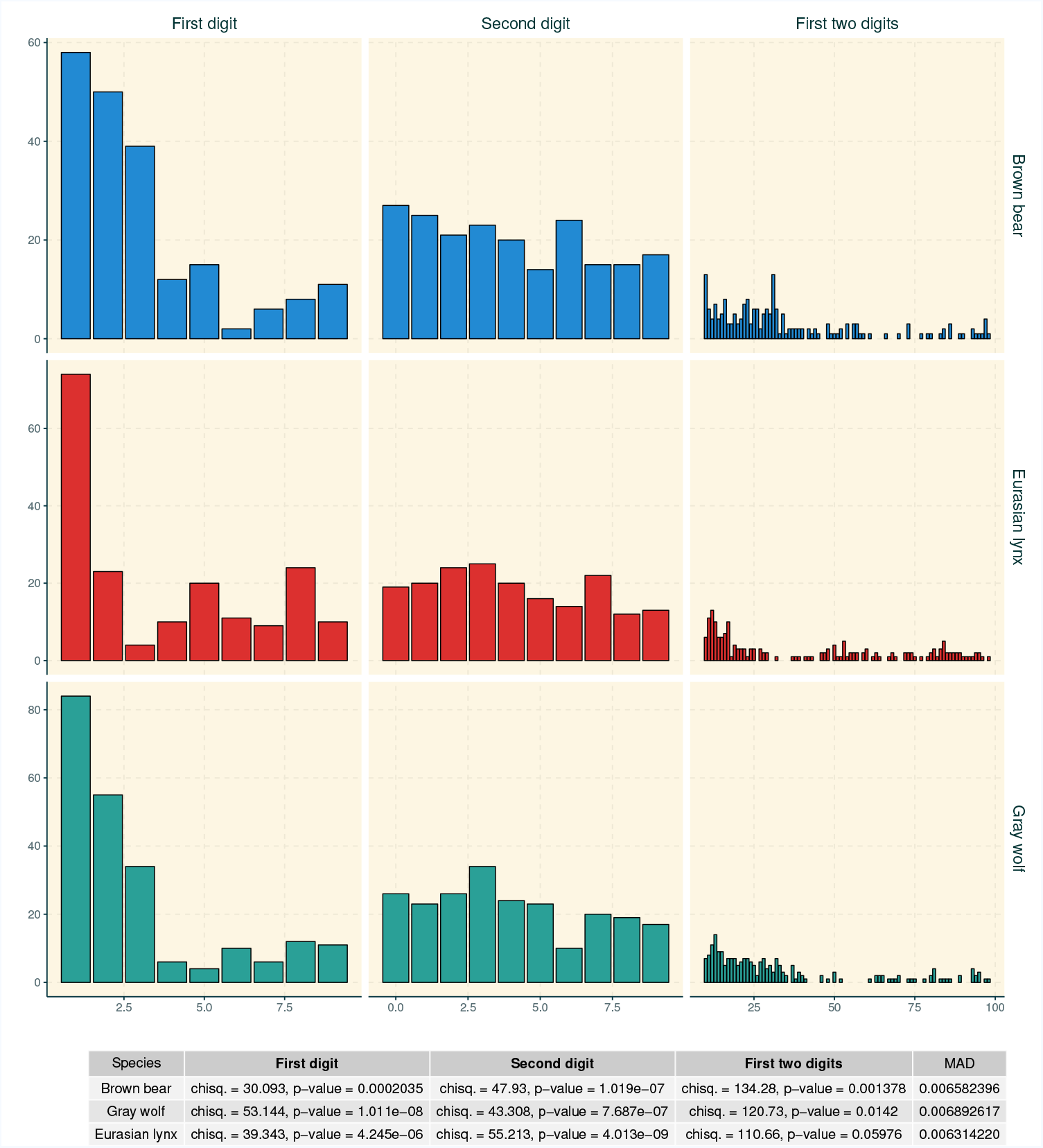
Frequency of the first, second and first two digits of regional population estimates of large carnivores in Romania (Popescu et al., 2016).

On the other hand, the scenario was more complex for whaling data from the former URSS fleet. While the chi-square test indicated that the first and second digit did not conform to a Benford’s distribution, and the MAD exceeded the cautionary threshold of 0.0044 suggested by Nigrini (2012) the chi-square test of the first couple of digits was non-significant, not deviating from a Benford’s distribution (Fig. 3).

**Figure 3: Fig.3.**
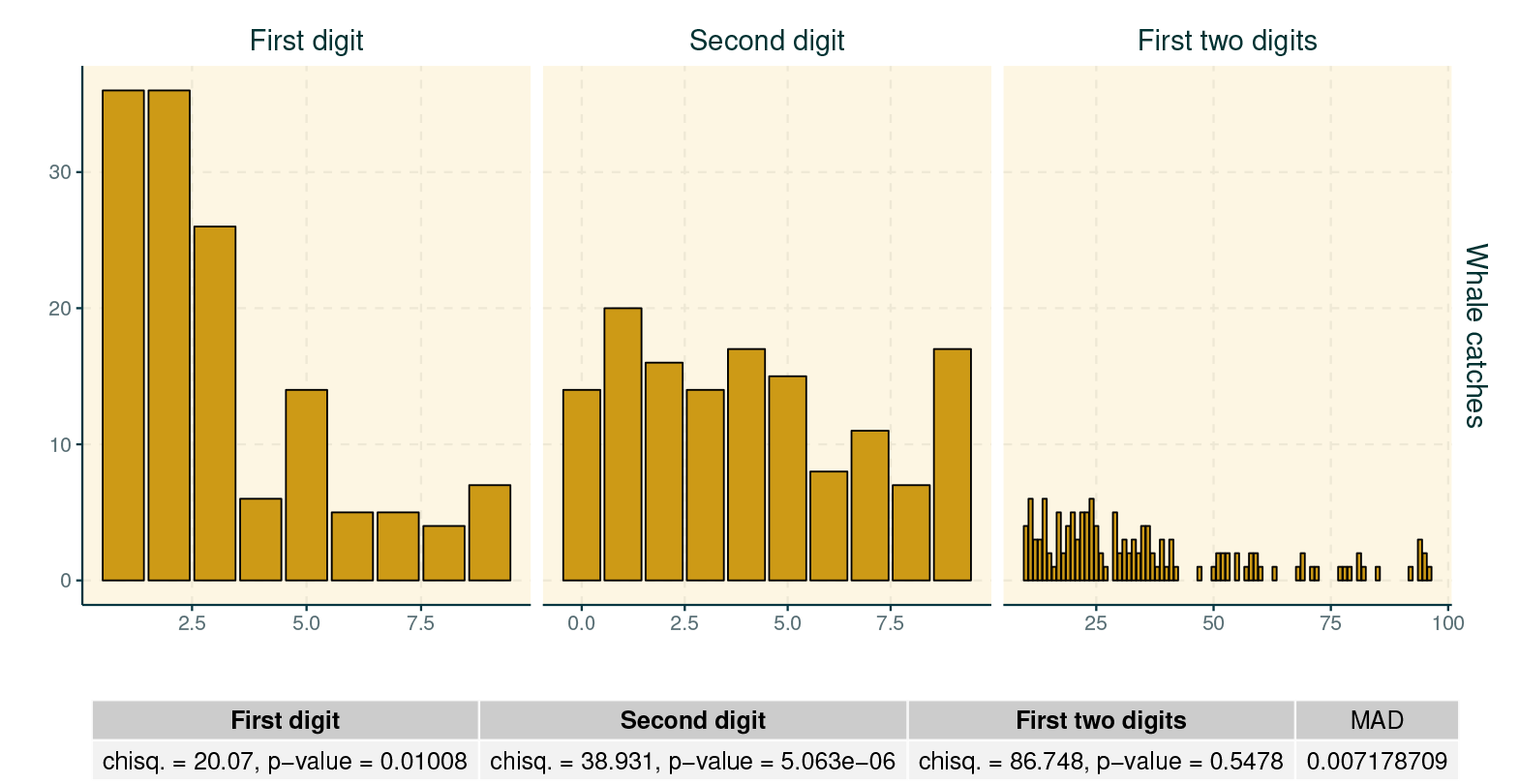
Frequency of the first, second and first two digits of Soviet whale catches in the Pacific Ocean (Clapham and Ivashchenko, 2009; Ivashchenko et al., 2013).

## Discussion

This validation study confirms the potential of digit-based tests based on the Benford’s law for auditing data in ecology and conservation. We believe that inspecting the frequency of the two leading digits of monitoring and harvesting data about natural resources will provide conservationist with the opportunity to detect anomalies that might underlie data manipulation or falsification. Then, efforts might be focused on these datasets, asking for supplementary information about data collection and weighting the evidence about data quality. In our research, the distribution of the first and second digits of two fraudolent numerical series, deviated from the expected Benford’s distribution that these digits should have followed, absent fraud. This was evident both for large carnivore population estimates from Popescu et al. (2016), where digits deviated for all the three species, and for commercial whale catches from Ivashchenko et al. (2013). On the other hand, the inspection of frequencies of the first couple of digits, taken together, was ambiguous: while goodness-of-fit testing and the MAD confirmed their anomalous distribution in the case of data about large carnivores in Romania, they were unable to detect falsification in whale catches from the Soviet fleet. Soviet data contained information about five different whale species, killed by four whaling fleets over a huge geographical scale: it is possible that falsification was not homogeneous across the species and fleets. Certainly it was heterogeneous across years, for the different whale species (Fig. 4).

**Figure 4: Fig.4.**
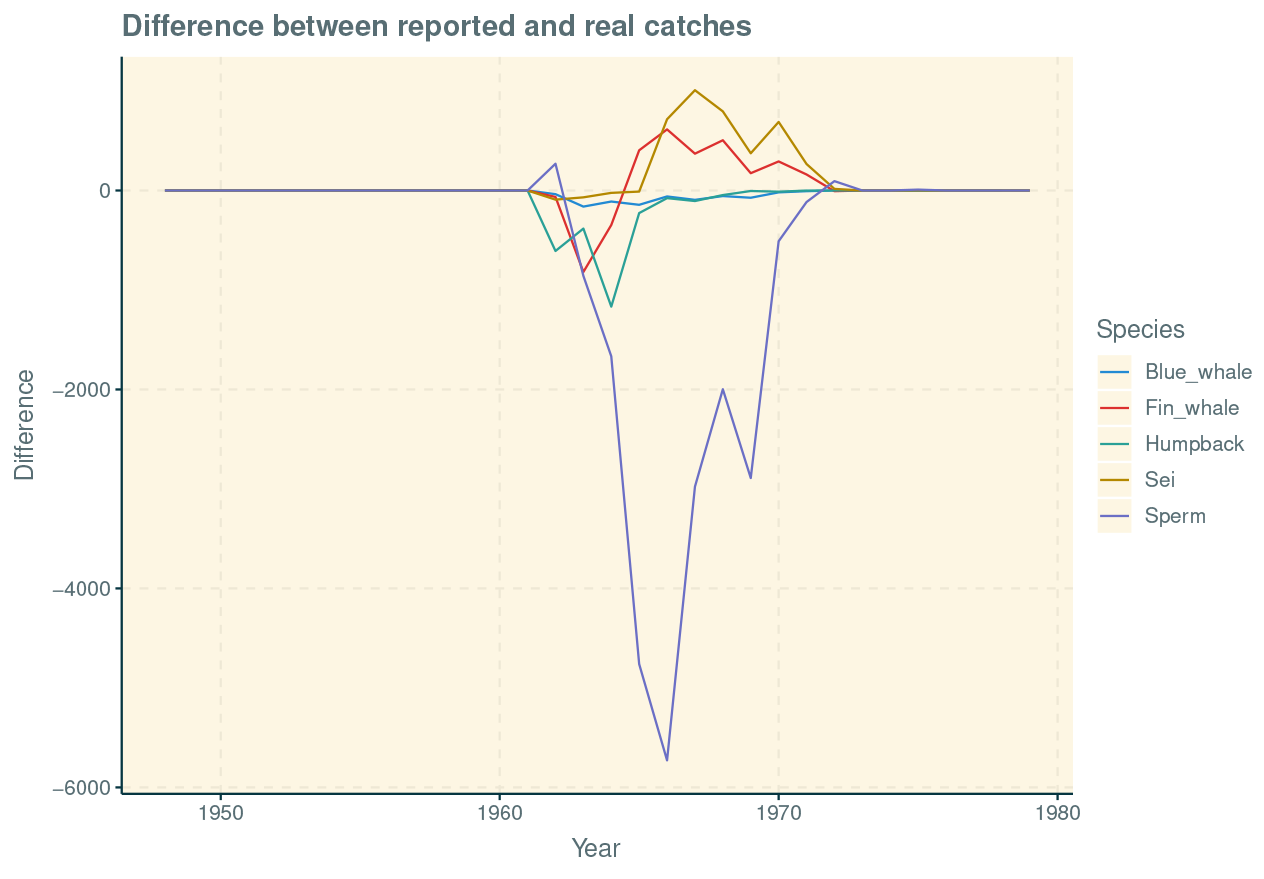
Differences in reported and real Soviet whale catches in the Pacific Ocean, across time (Clapham and Ivashchenko, 2009; Ivashchenko et al., 2013).

This inconsistency between two-digits and single-digit testing in whaling data delivers two messages. Firstly, it confirms the idea that multiple tests and approaches should be simultaneously adopted, to obtain a comprehensive picture about the data at hand (Nigrini, 2012), going beyond the superficial impression that would be obtained by using only one method. If we had adopted the chi-square goodness-of-fit test for the first two digits, alone, we would have oversimplified the interpretation of whale catches, that on the other hand showed an anomalous distribution of single digits and a very high value of the MAD. Secondly, it suggests to be cautious in using mixtures of very heterogeneous data for evaluating auditing systems as a whole, as we did for the Soviet whaling industry. Pooling together data collected from different sources aggregates information prone to different degrees and modes of falsification. For example, in our case study about Soviet whaling, the discrepancy between reported and real catches peaked between 1960s and the early 1970s, and it varied across fleets and species (Ivashchenko et al., 2013, Fig.4). It is plausible that this heterogeneity in the manipulation of data hampered the functioning of two-digits goodness-of-fit testing on the overall catches. We believe that future studies should compare the effectiveness of more refined goodness-of-fit tests (Barabesi et al., 2018; Lesperance et al., 2016) and the combination of these tests with other approaches, such as machine learning or the inspection of last-digits (Badal-Valero et al., 2018; Beber and Scacco, 2012) for detecting anomalies in mixtures of data. We also believe that other statistical approaches for the detection of manipulated data should be tested in ecology and conservation, because the Benford’s law has some precise distributional preconditions. These are often not respected, for example, by biometric or presence-absence data, which usually follow a Gaussian or a binomial distribution. Considered that some famous cases of data falsification in fisheries involved misreporting of length data to respect minimum sizes (Clapham and Ivashchenko, 2016, 2018), the development of approaches other than the Benford’s law should be a priority for conservationists. Because many biometric data for animals and plants are numeric series with four or more digits, statistical tests based the on expected uniformity of the last digits might be a valuable approach and future validation studies should test their potential.

However, from this research it must be clear that digit-based approaches, like the Benford’s distribution, are not meant to replace more refined techniques, such as machine learning or the comparison of data with complex and realistic population models (DiMininin 2018 a,b; Tulloch et al., 2018). Indeed, we believe that they could be very useful as early detection tools, like a warning light that lights up whenever the data do not look convincing. In the busy and messy field of ecology and conservation, warning lights might be very informative to detect problems as soon as possible and to avoid the contamination of decision making with wrong information. Their application would be similar to that of barcoding techniques in merceologic analysis, for auditing scams in food products or wood (Barcaccia et al., 2015; Godbout et al., 2018; Quinto et al., 2016). Once researchers decide to focus on a specific datasets that do not look trustworthy, they could opt for more in-depth and sophisticated approaches. Digit-based tests, in conjunction with other approaches such as specialized questioning techniques and structured methods for decision making (Nuno and St. John, 2015; Mukherjee et al., 2018), might become important tools for a more reliable and transparent conservation science.

